# Exploring Genomic Signatures of Selection in Guineafowl and Chicken Populations Across Diverse Climatic Zones: A Comparative Analysis of Africa, Asia, and Europe Research Article

**DOI:** 10.1101/2025.01.22.634375

**Authors:** Grace Moraa, Stephen Ogada, Stephen Kuria, Jacqueline Lichoti, Sheila Ommeh

**Affiliations:** Human Evolution, Department of Organismal Biology, Evolutionary Biology Centre, Uppsala University, Uppsala, Sweden; Institute for Biotechnology Research (IBR), Jomo Kenyatta University of Agriculture and Technology (JKUAT), P.O Box 62000, City Square 00200, Nairobi-Kenya; Central Veterinary Laboratories Kabete, State Department of Livestock, Ministry of Agriculture, Livestock and Fisheries, Private Bag, 00625, Nairobi-Kenya; Center for Animal Science, Queensland Alliance for Agriculture & Food Innovation, The University of Queensland, St Lucia QLD 4072

**Keywords:** genomic signatures, selection, climate change, environmental adaptation, guineafowl, chicken

## Abstract

This study aimed to investigate the signatures of selection in guineafowl and chicken genomes adapted to divergent climatic conditions across Africa, Asia, and Europe. We used whole genome sequence data of guineafowls and chicken from selected countries in these continents. To identify the signatures of selection, we employed three population genomics methods: F*_ST_,* integrated Haplotype Score (iHS), and Cross-Population Extended Haplotype Homozygosity (XP-EHH). Our findings revealed enriched terms related to metabolic processes, response to stimulus, signaling, and developmental processes, all of which play a role in the stress response for both guineafowls and chickens. Several candidate genes such as *CRYGN*, *BRAF*, *MAP3K2*, *ANGPT2*, *COL1A1*, *ATP13A4*, and *SLC66A1*, were among the positively selected candidate genes. Most of the candidate genes selected and the significant pathways identified play various roles in stress response in these two poultry species. Examination of divergent populations has provided new insights into genes potentially under selection for tolerance to the population’s indigenous environment, serving as a baseline for examining the genomic contributions to tolerance adaption. The knowledge gained from this research establishes a valuable foundation and guide for molecular breeding and conservation.

## 1. Introduction

Chicken (*Gallus gallus domesticus*), is an extensively distributed poultry species, and its initial domestication center is believed to be the Indus Valley in Southeast Asia (Peters et al., 2016; Zeuner, 1963). Studies have indicated that the Asian Red Jungle Fowl (*Gallus gallus*) is the ‘mother of all domesticated chicken’ and it was domesticated between 8000 to 5440 BC (Xiang et al., 2014). On the other hand, guineafowl (*Numida meleagris*) domestication has been traced to the African continent in Mali and Sudan about 2,000 years before present (BP) (Crawford, 1990; Gifford-Gonzalez & Hanotte, 2011; Larson & Fuller, 2014). Domestic helmeted guineafowls were domesticated from wild helmeted guineafowls. Additionally, there are other guineafowl species, including crested and vulturine, that are hunted in their natural wild habitats (Shen et al., 2021; Murunga et al., 2018). Since their domestication, chicken and guineafowls have dispersed to various parts of the world and they have developed into thousands of breeds with remarkable phenotypic characteristics in morphology, physiology, and behavior (Guo et al., 2016, 2022; Ulfah et al., 2016; Wang et al., 2020; Li et al., 2019; Moiseyeva et al., 2003). The wide range of chicken and guineafowl species is due to significant differences in resilience that have resulted from evolved genetic adaptations and artificial and natural selection (Thornton et al., 2009; Moraa et al., 2015). They have since adapted to their natural environments spanning the tropical, Mediterranean, and Arid climates in Africa, Monsoon, temperate, and Arid climates in Asia, and temperate and subarctic climates in Europe (Ulfah et al., 2016; Wang et al., 2015).

These two poultry species are among the most important sources of high-quality meat and eggs for the livelihood of rural communities across the world. Guineafowls are farmed in most African, Asian, and European countries as traditional village poultry. However, in most European countries like France, Italy, Belgium, Scandinavian countries, and the United States, they are intensively bred for commercial purposes (Araújo et al., 2023; Nahashon et al., 2006; Portillo-Salgado et al., 2022). In 2010, France produced 75% of European guineafowl production and 66% of the world’s production, and in 2017, it produced 30,000 tonnes, making it the leading guineafowl producer in the world (Vignal et al., 2019). The United States leads in poultry production at 23 million tonnes in 2019, followed by China at 20 million tonnes and Brazil at 16 million tonnes (FAO, 2022).

As the threat of climate change continues to grow, the adaptability of poultry species such as chicken and guineafowls becomes increasingly vital (Blench & MacDonald, 2006; Portillo-Salgado et al., 2022). These species have undergone long-term natural selection, which has led to favorable changes in their allele frequencies, allowing them to thrive in their respective natural environmental conditions (Asadollahpour et al., 2022). However, as environmental conditions shift due to climate change, it remains to be seen if these species’ adaptations will be sufficient to ensure their survival in the long run. Therefore, uncovering the genetic basis underlying heat stress response has both theoretical importance and practical applications in facilitating breeding programs to address heat stress (Guo et al., 2022).

Selection signatures are distinct genetic patterns or footprints left behind in the genomic regions subjected to selection (Saravanan et al., 2020). Over time, selection pressures affect genome structure and leave signatures in specific regions of the genome, such as increased allele frequencies, extensive linkage disequilibrium, homozygous genotypes, and decreased local diversity (Nielsen, 2005; Qanbari & Simianer, 2014; Rostamzadeh et al., 2021; Sabeti et al., 2006). The study and identification of loci under natural selection is of great interest in various research fields, as it could help researchers better understand the genetic mechanism underlying natural selection and adaptability to various environments (Asadollahpour et al., 2022; Foll & Gaggiotti, 2008). Signatures of selection found in animal genomes have led to the identification of genes that could be used to breed animals with preferred traits (Bian et al., 2019). For instance, the *BMP2*, *SP3*, *PROKR1* genes, which play an important role in fat tail formation, could be a guide for molecular breeding and conservation in Chinese indigenous sheep (Yuan et al., 2017). The *PICALM* gene associated with αS1-casein content in cattle milk has the potential to be used in breeding programs to improve the productivity of dromedary camels through genomic selection (Bahbahani et al., 2019).

To decipher the genetic mechanisms that are involved in the domestication, adaptation, and phenotypic differentiation of individuals belonging to the same species, the detection of signatures of selection using different statistics has been performed in various species, including chicken (Almeida et al., 2019; Fleming et al., 2017; Lawal et al., 2018; Wang et al., 2020), cattle (Jahuey-Martínez et al., 2019; Yurchenko et al., 2018; Hozé et al., 2014; Randhawa et al., 2014), sheep (Liu et al., 2016; Wang et al., 2019), pigs (Zhang et al., 2019), and goats (Bertolini et al., 2018; Onzima et al., 2018).

Despite the economic importance of chicken and guineafowls, the genomics of adaptation, especially in the guineafowls remains an enigma (Mahammi et al., 2016). Hence, this study employed WGS data of diverse chicken and guineafowl ecotypes from selected countries in Africa, Asia, and Europe to identify genomic regions and potential candidate genes that may be subject to selective pressure for adaptation to varying environments in the face of climate change. This understanding is significant for refining approaches to develop climate-resilient poultry breeds that can be effectively utilized and preserved.

## 2. Materials and Methods

### 2.1. Dataset

We downloaded 72 guineafowl and 14 chicken whole-genome sequence data previously published from the Sequence Read Archive (SRA) in the NCBI database. The guineafowls study accession numbers (SRP267755 and SRP166710) included samples from Europe (five France, five Italy, seven Hungary) Africa (five Burkina Faso, one Benin, one South Africa, 16 Kenya, 10 Sudan, 10 Nigeria), Asia (six China, six Iran). Out of the 72 downloaded guineafowl genomes, five were from the vulturine guineafowl species from Kenya, one was from the crested guineafowl species from Kenya, and 64 were from the wild and domestic helmeted guineafowl species from various countries in Africa, Asia, and Europe. The guineafowl genome data were from experimental populations that had been genotyped and described in previous studies (Shen et al., 2021; Vignal et al., 2019) (Table S1).

The expansion of trade and exploration enabled the dispersion of helmeted guineafowls from their centers of domestication in Africa to various parts of the world, where they have since adapted to the local environmental conditions (Larson & Fuller, 2014; Vignal et al., 2019). Previous studies on genetic diversity, utilizing mitochondrial DNA (Murunga et al., 2018; Adeola et al., 2015) and microsatellite markers (Botchway et al., 2013; Kayang et al., 2010), revealed no significant genetic structuring among African domestic helmeted guineafowl populations. This suggests a recent domestication followed by a rapid and extensive dispersal across Africa.

On the other hand, the chicken dataset accession numbers (SRP142580 and SRP040477) included chicken from Africa (nine Ethiopia) (Lawal et al., 2018), and Asia (five China) (Wang et al., 2015) (Table S2). The chicken whole genome data downloaded originated from indigenous domestic village chickens from Horro and Jarso regions in Ethiopia (Lawal et al., 2018). In China, the whole genome sequence data was from the red junglefowl from the regions of Yunnan and Hainan (Wang et al., 2015). The chicken data were from experimental populations that had been genotyped and described in previous studies (Lawal et al., 2018; Wang et al., 2015).

### 2.2. Whole-Genome Sequence Quality Control Processing and Variant Calling

Trimmomatic v0.36 (Bolger et al., 2014) was used to filter out low-quality reads and adapter sequences of raw data, based on default parameters. The guineafowl paired-end reads were mapped to the guineafowl reference genome (NumMel1.0), while the chicken paired-end reads were mapped to the chicken reference genome (GRCg6a), from the NCBI database. Each individual was aligned using Burrows-Wheeler Aligner (BWA) with the option “BWA-MEM” algorithm with conventional parameters (Li & Durbin, 2010). We followed the recommendations of the Broad Institute Genome Analysis Toolkit v.4.1.3.0 (GATK) Best Practices for the preprocessing workflow preceding variant discovery (McKenna et al., 2010). Picard tools v.1.56 (Li & Durbin, 2010) was used to sort and merge the alignment files by coordinates, index them, calculate the alignment matrices, and to MarkDuplicates. Calling of variants for each sample was performed using the default parameters of the GATK “HaplotypeCaller”. Joint genotyping (GenotypeGVCFs) was done to identify variants simultaneously in all guineafowl and chicken samples. Single Nucleotide Polymorphisms (SNPs) and insertion/deletions (InDels) in the guineafowl and chicken genomes were filtered into two different files using GATK VariantFiltration. Hard filtering was performed to reduce false positive variants. To exclude SNP calling errors caused by incorrect mapping, only high-quality SNPs filtered by the VariantFiltration of GATK with options -filter "QD < 2.0" --filter-name "QD2" -filter "QUAL < 30.0" --filter-name "QUAL30" -filter "SOR > 4.0" -- filter-name "SOR3" -filter "FS > 60.0" --filter-name "FS60" -filter "MQ < 40.0" --filter-name "MQ40" -filter "MQRankSum < -12.5" --filter-name "MQRankSum-12.5" -filter "ReadPosRankSum < -8.0" --filter-name "ReadPosRankSum-8" were retained for subsequent analyses (Liu et al., 2022). To estimate SNP statistics like the number of total and average bi-allelic autosomal SNPs and the ratio of heterozygous and homozygous SNPs BCFtools v.1.10.2 was used.

### 2.3. Population Structure Analysis

Population structure was inferred by Principal component analysis (PCA), and ADMIXTURE analysis for the guineafowl and chicken populations. To minimize SNP redundancy, we pruned the two datasets using PLINK v1.90 (Purcell et al., 2007) with options “-indep-pairwise 50 10 0.2”(Anderson et al., 2010). Based on pruned SNPs, PCA was performed with smartpca in EIGENSOFT v7.2.0 (Patterson et al., 2006). Then, by using the first (PC1) and second (PC2) principal components, the figures were plotted with R v4.0.5 (Lüdecke et al., 2021) using an in-house script. The unsupervised hierarchical clustering was performed for all guineafowl and chicken populations using ADMIXTURE v1.3.0 (Alexander et al., 2009). To identify the best value of K clusters, we ran ADMIXTURE with cross-validation for values of K from 2 to 6 for guineafowls and 2 to 5 for chicken.

### 2.4. Selective Sweep Analysis

To identify the candidate regions under positive selection in the various guineafowl and chicken populations, we calculated the fixation statistics (F*_ST_*) as previously described (Li et al., 2013). Differentiation between the combination of chicken populations from Ethiopia vs. China was evaluated. Differentiation in the guineafowl populations was evaluated for the following combinations: (1) Africa vs. Asia, (2) Africa vs. Europe, and (3) Asia vs. Europe. We estimated the F*_ST_* values by use of the VCFtools v0.1.13 (Holsinger & Weir, 2009), in a 500-kb sliding window with a 250-kb step size (Li et al., 2013; Qin et al., 2020). We empirically selected the top 0.1% F*_ST_* values as potential candidate regions under selection (Qin et al., 2020).

We also used two complementary EHH-derived methods, the Cross Population Extended Haplotype Homozygosity (XP-EHH) (Sabeti et al., 2007), and the Integrated Haplotype Score (iHS) (Voight et al., 2006), to detect signatures of selection. We used the XP-EHH as a cross-population method to detect signals of divergent selection between Ethiopia vs. China chicken populations. The following combinations of guineafowl populations were evaluated for the XP-EHH score: (1) Africa vs. Asia, (2) Africa vs. Europe, and (3) Asia vs. Europe. The iHS was used as a within-population method for the identification of signals of ongoing positive selection. We calculated the iHS values for the chicken populations from Ethiopia, and China, and guineafowl populations from Africa, Asia, and Europe. The guineafowl and chicken SNPs were phased using SHAPEIT 4.2.1 (Browning et al., 2021). The iHS and XP-EHH scores were calculated by rehh v3.2.2 (Gautier & Vitalis, 2012) with default parameters. To detect positive selection in guineafowl and chicken populations, the average XP-EHH and iHS scores were computed for 100-kb regions with a 50-kb overlap.

### 2.5. Functional Annotation for Candidate Genes

To uncover the biological functions of candidate genes, we performed the Gene Ontology (GO) Biological processes, Reactome Gene sets, CORUM, Canonical Pathways, and the Kyoto Encyclopedia of Genes and Genomes (KEGG) enrichment analysis on the Metascape website (http://metascape.org) (Kanehisa et al., 2019; Zhou et al., 2019). We used the single list analysis method in our annotations using the guineafowl and chicken reference genomes to assign genes to the corresponding terms. Enrichment tests were performed using the Benjamini-Hochberg *p*-value correction algorithm as described in Metascape (Zhou et al., 2019).

## 3. Results and Discussion

### 3.1. Population Structure Analysis in Guineafowls

The principal component analysis (PCA) supports the population relationships based on the average genetic distance. The plot of the first (PC1) and second (PC2) eigenvectors in guineafowls explained about 24.5% and 12.3% of the proportion of variations, respectively. Principal components (PCs) completely separated the three guineafowl populations according to their sampled locations and species. The vulturine guineafowls from Africa formed their cluster. Walker et al. (2004) and Vignal et al. (2019) both reported comparable findings, indicating a distinct separation between wild and domestic guineafowl populations. Most of the helmeted guineafowls from Africa were in the same cluster as those from Asia, and Europe. The crested guineafowl from Africa formed its cluster (Figure 1(a)). The results of the ADMIXTURE corroborated the finding of PCA. When K=6, there was no admixture in the crested guineafowl species from Africa, and in some helmeted guineafowl species from Africa. Four vulturine guineafowl species from Africa showed no admixture across all the K values (Figure 1(b)).

**Figure 1.**
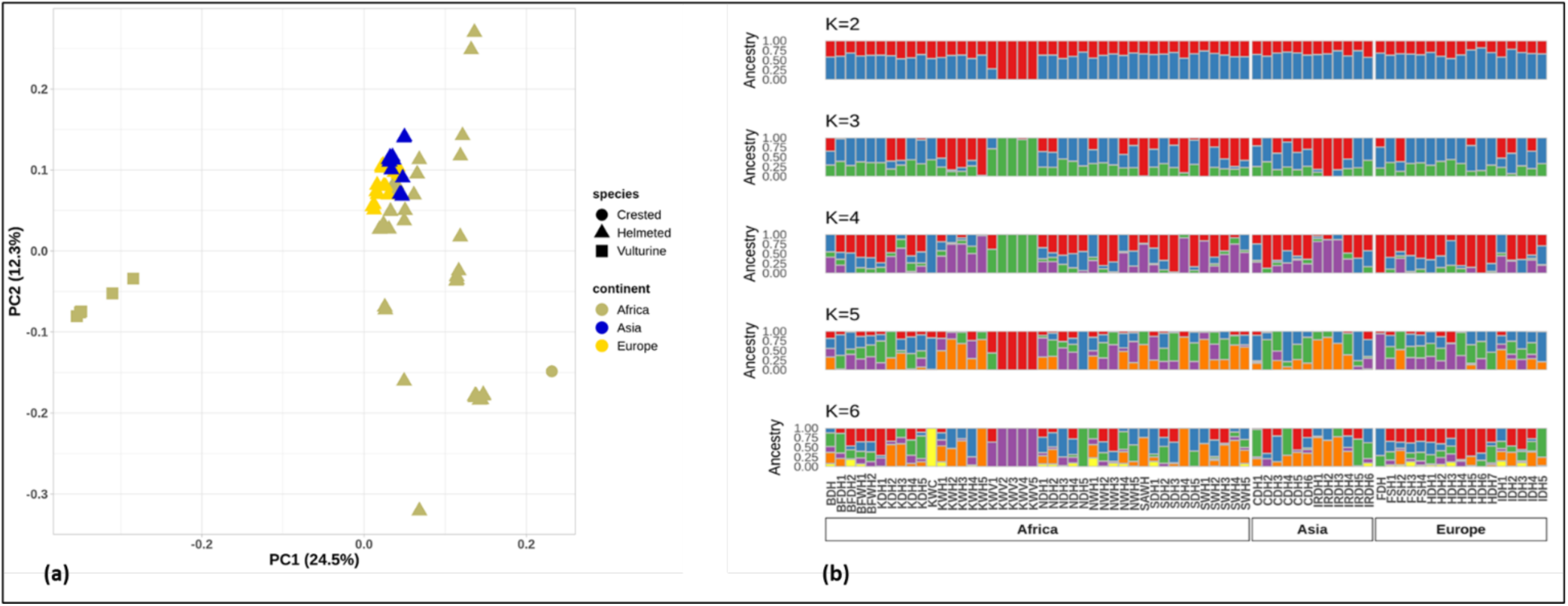
Genetic structure of guineafowls defined with PCA and ADMIXTURE analyses. (a) Guineafowl population structure as revealed by PCA. The variance explained by each PC is indicated within brackets. (b) Guineafowl population structure analysis as revealed by ADMIXTURE analysis. These results with inferred numbers of clusters from K=2 to K=6. The white lines separate the experimental samples that were from Africa, Asia, and Europe. Each sample is represented by a vertical bar.

Both PCA and ADMIXTURE analysis confirmed the genetic similarity between the domestic guineafowl populations from Africa, Asia, and Europe. However, the vulturine guineafowl species which is a wild guineafowl from Africa, showed a lack of admixture revealing a highly homogenous genetic composition. The admixture seen in the domestic helmeted guineafowls could be a result of the crossbreeding practices by farmers to obtain desired physical characteristics.

### 3.2. Population Structure Analysis in Chicken

The plot of the first (PC1) and second (PC2) eigenvectors explained about 16% and 13.1% of the proportion of variations, respectively. The PCs separated the chicken populations into three clusters; cluster 1 represents chicken from Yunnan (CYR1-CYR4) and Hainan (CHR1) in China, cluster 2 represents chicken from Jarso in Ethiopia (EJ1-EJ4), and cluster 3 represents chicken from Horro in Ethiopia (EH1-EH5) (Figure 2(a)). The results of the ADMIXTURE corroborated the finding of PCA, since at K=5, only one chicken sample from Jarso in Ethiopia (EJ4) showed admixture. Three chicken samples from Jarso in Ethiopia (EJ1-EJ3), and three chicken samples from Yunan in China (CYR1, CYR3, and CYR4) showed no admixture across all the K values (Figure 2(b)), indicating the genetic integrity and purity of these chicken breeds.

**Figure 2.**
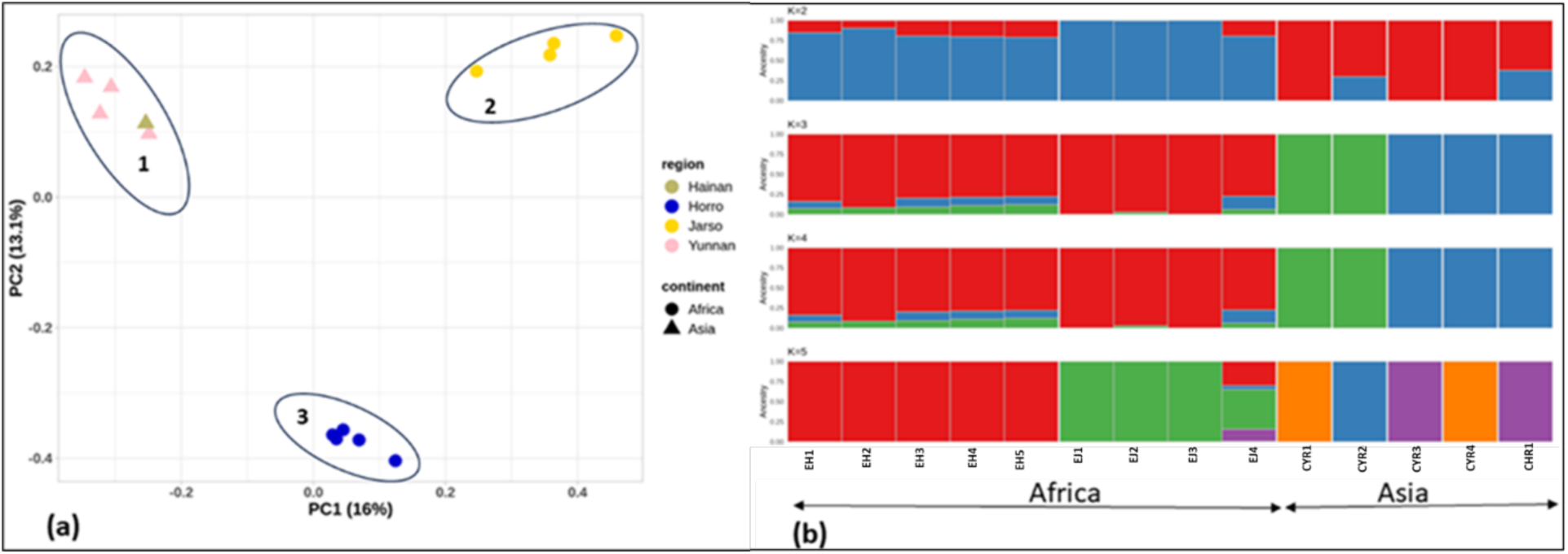
Genetic structure of chicken defined with PCA and ADMIXTURE analyses. (a) Chicken population structure as revealed by PCA. The variance explained by each PC is indicated within brackets. (b) Chicken population structure analysis as revealed by admixture analysis. These results with inferred numbers of clusters from K=2 to K=5. The white lines separate the experimental samples that were from Africa and Asia. Each sample is represented by a vertical bar.

### 3.3. Selection Sweep Signals in Guineafowls

#### 3.3.1. Selection Signatures Common in Guineafowl Populations from Africa and Asia

Some of the selection sweep regions were detected on chromosome 1 (NumMel1.0 position 57,687,262-57,756,865 bp), and chromosome 5 (NumMel1.0 position 57,878,739- 57,894,933 bp; 57,850,727-57,871,807 bp; 57,746,524-57,790,117 bp) by the F*_ST_* method (Figure 3(a)) and on chromosomes 2 (NumMel1.0 position 89,102,231-89,170,488 bp), chromosome 3 (NumMel1.0 position 89,063,612-89,108,133 bp; 88,662,013-88,692,902 bp), and chromosome 4 (NumMel1.0 position 88,828,418-88,902,944 bp) by the XP-EHH method (Figure 3(b)). Some of the significantly overrepresented GO terms and pathways include (1) response to stimulus category (GO:0009314, response to radiation), (2) signaling category (WP422, MAPK cascade), and, (3) metabolic process category (HSA-8978868, fatty acid metabolism) (Table S3, Figure S1A). The enriched terms could play several roles in the adaptation of the guineafowls to their varying environments. For instance, a metabolic process such as lipid metabolism, has been shown to play a role in the growth and reproduction performance of poultry (Wang et al., 2020). The primary role of fat cells is to store and release energy. This function is crucial for animals living in stressed environments as it allows them to regulate fat metabolism and maintain energy balance, facilitating adaptation to their surroundings (Guo et al., 2016; Zampiga et al., 2018; Wang et al., 2020). Response to radiation, which may include ultraviolet and ionizing radiation, could enhance the adaptation of guineafowls to high-intensity solar radiation in heat-stressed environments and during periods of heat waves.

**Figure 3.**
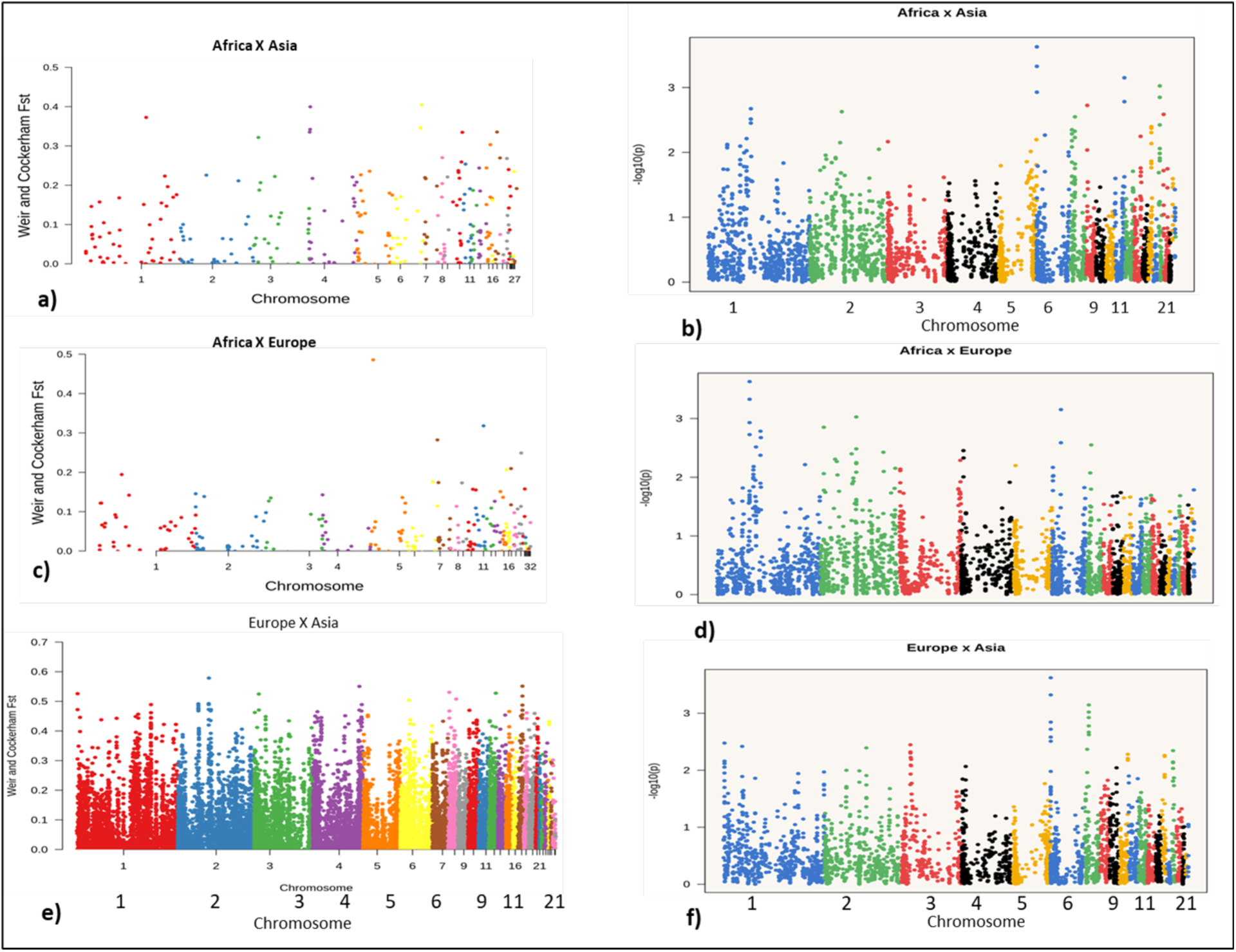
Distribution of F*_ST_* and XP-EHH scores amongst guineafowl populations from Africa, Asia and Europe. (a) Distribution of F*_ST_* scores comparing amongst Africa vs. Asia guineafowl populations. (b) Distribution of XP-EHH scores comparing amongst Africa vs. Asia guineafowl populations. (c) Distribution of F*_ST_* scores comparing amongst Africa vs. Europe guineafowl populations. (d) Distribution of XP-EHH scores comparing amongst Africa vs. Europe guineafowl populations. (e) Distribution of F*_ST_* scores comparing amongst Europe vs. Asia guineafowl populations. (f) Distribution of XP-EHH scores comparing amongst Europe vs. Asia guineafowl populations.

The response to radiation GO term was found to be enriched in the Dulong chicken, adapted to survive in low-temperature environments. Response to radiation is crucial for the survival of livestock in stressful environments (Wang et al., 2020). Some of the candidate genes that were selected for these pathways include *BRAF*, *ZNF236*, *GCLC*, *ANGPT2*, *MAP3K2*, *ERCC3*, *XRCC5*, and *HTT* (Table S3). The *MAP3K2* gene belongs to the mitogen-activated protein kinase family, which transmits signals from extracellular stimuli to the nucleus (Guo et al., 2020). Earlier studies have indicated that this particular gene is significantly involved in controlling a range of biological functions in different animal species. These functions include cellular growth, cellular specialization, reproductive processes, immune system activity, secretion, and stress response (Wang et al., 2020; Guo et al., 2020; Sheng et al., 2020). The *BRAF* gene has been shown to mediate the *SLIT2-SRGAP1-CDC42*-induced granulosa cell proliferation and differentiation in chicken (Shen et al., 2022). Hence, this particular gene may play a crucial role in the maturation of granulosa cells and the process of ovulation, thereby impacting the reproductive capabilities and egg-laying performance of guineafowls inhabiting heat-stressed environments.

#### 3.3.2. Selection Signatures Common in Guineafowl Populations from Africa and Europe

Selection sweeps were observed on chromosome 2 (NumMel1.0 position 5,510,580- 5,515,600 bp), chromosome 4 (NumMel1.0 position 4,837,773-4,841,119 bp), chromosome 16 (NumMel1.0 position 4,816,405-4,822,410 bp), chromosome 20 (NumMel1.0 position 4,553,693-4,565,139 bp), chromosome 21 (NumMel1.0 position 4,664,120-4,671,719 bp), and chromosome 25 (NumMel1.0 position 5,413,615-5,440,266 bp) by the F*_ST_* method (Figure 3(c)) and on chromosome 4 (NumMel1.0 position 12,463,584-12,474,248 bp), and chromosome 8 (NumMel1.0 position 12,439,902-12,461,344 bp) by the XP-EHH method (Figure 3(d)). Candidate genes located in the selected regions were subjected to GO analysis (Table S4). Some of the top-level enriched terms were involved in (1) the developmental process category (GO:0021761, limbic system development), and, (2) the metabolic process category (GO:0016042, lipid catabolic process) (Figure S1B).

The lipid catabolic process is crucial for the growth and reproduction of animals in stressed environments. In poultry, embryo growth depends on lipid metabolism sourced from the diet or produced in the liver. These lipids, transported in the bloodstream as small molecules, reach the ovaries and are stored in the yolk. Serving as an energy source, the yolk provides the embryo with the necessary energy for development (Wang et al., 2020). Heat stress significantly impacts embryo development in poultry. Given that guineafowls are primarily raised for meat and eggs, the lipid catabolic process plays a vital role in enhancing embryo development and hatchability in stressed environments.

The limbic system development also plays a role during stress response. When confronted with stress, specific neural pathways in the limbic system and prefrontal cortex of the brain are activated, leading to the release of adrenaline and noradrenaline. The stress hormones in the limbic system appear to be significant in the adaptation to sudden stressors like heat stress (Durosaro et al., 2023), and this could be important in the adaptation of guineafowl populations to stress environments.

The *CRYGN*, *SLC16A2*, *SLC20A1*, *SLC16A14*, *SLC27A4*, *LRRC8A*, *HNF4A*, *CNBP*, *SREBF1*, *GHSR*, *ATP2B4*, *ATP13A2*, and *ADA* genes were some of the candidate genes selected in enriched pathways (Table S4). The *SLC* family of genes which were involved in several pathways has been shown to facilitate the transfer of molecules like sugars, nucleotides, and amino acids to maintain a constant internal environment (Shi et al., 2022; Mueckler & Thorens, 2013). Several other genes within the *SLC* family have been proposed as having high potential as heat stress biomarkers for different organisms (Bao et al., 2017; Shi et al., 2022; Yadav et al., 2018). The *SLC16A12* was among the positively selected genes in diverse breeds of chicken in China, and it has been suggested to be one of the main genetic contributors to chicken domestication (Li et al., 2019). Hence, the *SLC* gene family, in collaboration with other genes that were positively selected, could play a significant role in the adaptation of guineafowls to diverse environmental conditions.

#### 3.3.3. Selection Signatures Common in Guineafowl Populations from Europe and Asia

Strong putative selection signature regions were detected on chromosome 1 (NumMel1.0 position 59,730,381-59,765,151 bp), chromosome 2 (NumMel1.0 position 59,880,060-59,932,870 bp), chromosome 4 (NumMel1.0 position 59,367,873-59,373,467 bp; 59,344,326-59,365,288 bp), and chromosome 6 (NumMel1.0 position 59,184,416- 59,184,943 bp; 60,067,955-60,083,936 bp) by the F*_ST_* method (Figure 3(e)) and on chromosome 5 (NumMel1.0 position 18,091,305-18,114,688 bp), chromosome 9 (NumMel1.0 position 18,192,774-18,225,307 bp), and chromosome 10 (NumMel1.0 position 18,232,994-18,239,840 bp) by the XP-EHH method (Figure 3(f)). Several GO terms were enriched (Table S5). One of the significantly enriched terms was the Developmental process category (GO:0060428, lung epithelium development), and (GO:0007420, brain development) (Figure S1C).

The enrichment of developmental processes indicates the guineafowls’ ability to flourish in varying environments. This term was also found to be enriched in chickens adapted to high-altitude and high-humidity environments (Wang et al., 2020). The process of lung epithelium development may provide a mechanism for guineafowls to enhance the transportation and utilization of oxygen, thereby facilitating their adaptation to specific environments.

Some of the candidate genes that were identified include *ARF6*, *BMP4*, *ALOX5*, *DNM1L*, *EXOSC9*, *NUP153*, *CDT1*, *MAP2K1*, and *BBS7* (Table S5). The *MAP2K1* gene belongs to the mitogen-activated protein kinase family, and it is involved in the transmission of signals from extracellular stimuli to the nucleus (Guo et al., 2020). This gene has also been shown to play a role in regulating physiological processes like cell proliferation, cell differentiation, reproduction, and stress response in various livestock species (Wang et al., 2020; Guo et al., 2020; Sheng et al., 2020).

#### 3.3.4. Selection Signatures Common in Guineafowl Populations from Africa

Strong selection sweeps were observed on chromosome 1 (Figure 4(a)). Candidate genes located in the selected regions were used for the detection of enriched GO terms. Some of the significant GO terms include (1) developmental process category (GO:0051963, regulation of synapse assembly), (GO:0045664, regulation of neuron differentiation), (2) response to stimulus category (GO:0080135, regulation of cellular response to stress), and, signaling category (GO:1905114, cell surface receptor signaling pathway involved in cell-cell signaling) (Figure S2A). The *BAK1*, *XDH*, *TNFSF10*, and *PLCG1* genes were some of the candidate genes that were involved in the enriched pathways (Table S6).

**Figure 4.**
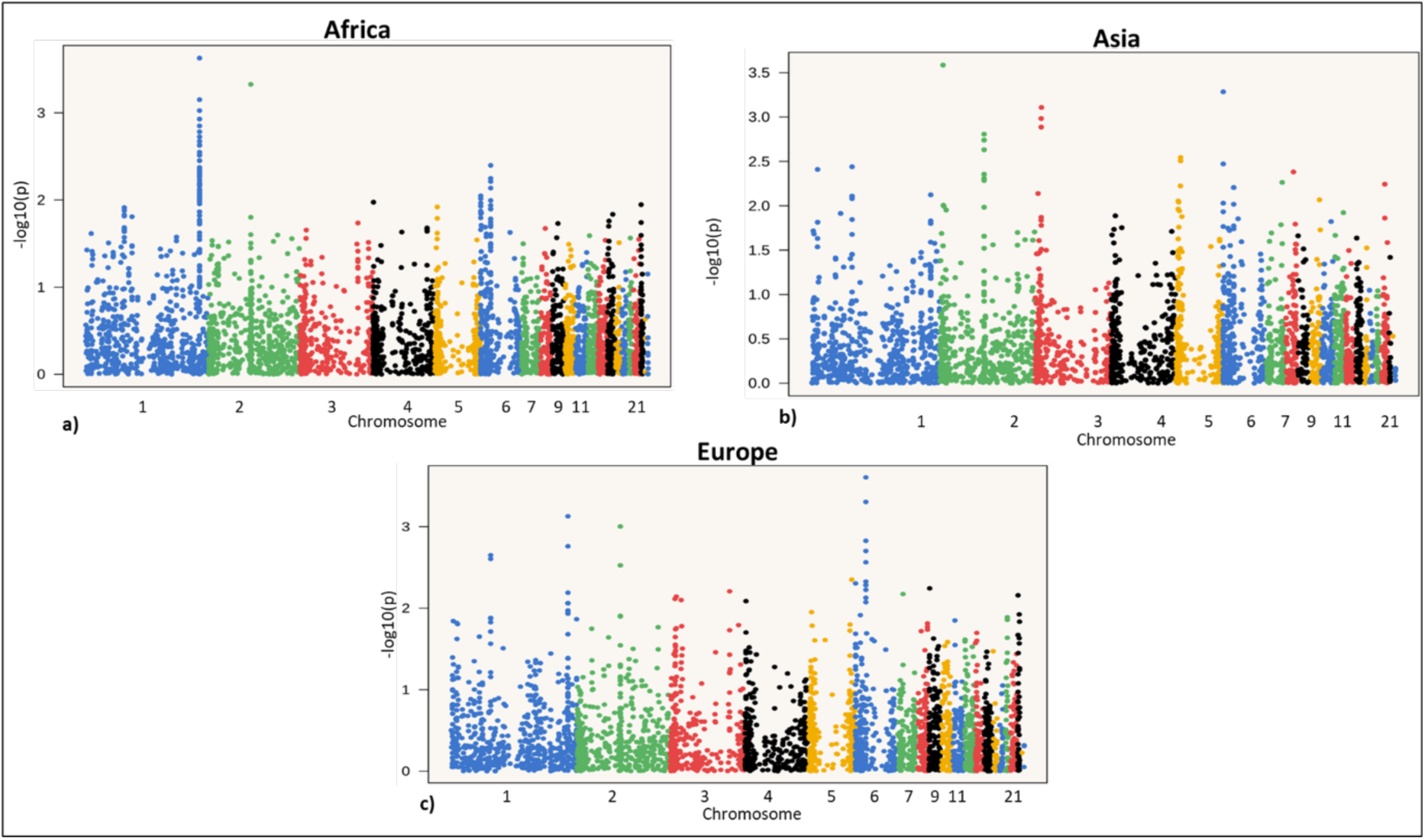
Genome-wide distribution of iHS values for the guineafowl populations from Africa, Asia, and Europe. (a) Distribution of iHS values for the guineafowl populations from Africa. (b) Distribution of iHS values for the guineafowl populations from Asia. (c) Distribution of iHS values for the guineafowl populations from Europe

Studies have shown that several genes are associated with changes in behavior related to domestication, such as decreased fear, increased exploration, altered learning, and memory capacity (Forrest et al., 2018; Schafe et al., 2001; Sheng & Kim, 2002; Sweatt, 2004). These changes are considered regulated by neurotransmission and signal transduction that affect synaptic plasticity and neural circuits involved in emotional, social, and cognitive functions (Forrest et al., 2018; Schafe et al., 2001; Sheng & Kim, 2002; Sweatt, 2004). Therefore, the developmental process of synapse assembly and regulation of neuron differentiation, which were enriched in guineafowl populations from Africa could play an important role in the domestication of guineafowls, and in their adaptation to different environments.

#### 3.3.5. Selection Signatures Common in Guineafowl Populations from Asia

The strongest putative regions of selection were on chromosome 13 (NumMel1.0 position 4,732,348-4,751,869 bp), chromosome 16 (NumMel1.0 position 5,154,885-5,160,043 bp), chromosome 19 (NumMel1.0 position 4,879,891-4,895,599 bp), and chromosome 20 (4,820,225-4,823,211 bp; 5,045,127-5,052,041bp) (Figure 4(b)). The selected genes were subjected to GO and some of the significant terms included (1) localization category (GO:0051235, maintenance of location), (2) response to stimulus category (GO:0071453, cellular response to oxygen levels), and (3) biological regulation category (R-HAS-556833, metabolism of lipids) (Figure S2B). The enrichment of the cellular response to oxygen levels GO term is significant, as guineafowls have evolved mechanisms to cope with varying oxygen levels in their habitats. For example, high-altitude environments often have lower oxygen levels. Guineafowls inhabiting these environments might exhibit specific cellular responses to ensure optimal oxygen utilization (Wang et al., 2020). Some of the candidate genes that were identified included the *SLC66A1*, *SLC27A4*, *LRRC8A*, *CNBP*, *SREBF1*, *HNF4A*, *BRCA1*, *PINK1*, *MUL1*, *MAP2K3*, and *SDC4* (Table S7). The *SLC* family of genes, which were also selected in the guineafowl populations from Africa and Europe, have been shown to play various roles in homeostasis (Shi et al., 2022; Mueckler & Thorens, 2013) and in stress regulation in different organisms (Bao et al., 2017; Shi et al., 2022; Yadav et al., 2018). One of the genes in the *SLC* family of genes, *SLC41A2*, has previously been identified as being under selection in guineafowl populations in a region that was selected after importation into Europe. It was suggested to be related to selection after the domestication of the guineafowls (Vignal et al., 2019)

#### 3.3.6. Selection Signatures Common in Guineafowl Populations from Europe

Figure 4(c) shows the genome-wide distribution of iHS values across the genome of the guineafowl population from Europe. Selection sweeps were detected on chromosome 3 (NumMel1.0 position 15,743,601-15,760,242 bp), chromosome 4 (NumMel1.0 position 15,543,048-15,575,141 bp; 15,506,128-15,541,415 bp; 15,790,768-15,933,899 bp), chromosome 5 (NumMel1.0 position 15,525,897-15,648,603 bp), and chromosome 10 (NumMel1.0 position 15,673,441-16,117,582 bp). Several candidate genes were selected and when subjected to GO analysis, 39 enriched terms were identified. The homeostatic process category (GO:0006874, cellular calcium ion homeostasis) was one of the significant GO terms (Figure S2C). *ATP13A4*, *ATP13A5*, *CDH13*, *DOCK10*, *PLCE1*, and *POLR1B*, were identified as potential candidate genes for stress adaptation (Table S8).

The *ATP13A4* and *ATP13A5* genes were identified in the cellular calcium ion homeostasis pathway. These genes belong to the large family of genes involved in ATP-dependent ion transport involved in calcium homeostasis (Sim & Park, 2023; Szigeti & Kellermayer, 2006). Maintenance of calcium ion homeostasis has been reported to be related to temperate environment adaptation in previous studies in chickens from highland areas of China (Nan et al., 2023; Zhang et al., 2016; Wang et al., 2015). These genes, therefore, play an important role in the adaptation of guineafowls to various climates in their habitats.

### 3.4. Selection Sweep Signals in Chicken

#### 3.4.1. Selection Signatures Common in Chicken Populations from Africa

Strong selective sweeps were observed on chromosome 1 (GRCg6a position 146,060,187- 146,065,582 bp; 146,022,451-146,033,636 bp), and chromosome 2 (GRCg6a position 146,163,339-146,186,941 bp; 145,987,868-146,024,793 bp) (Figure 5(a)). The top-level GO enriched terms were involved in signaling categories (GO:0046578, regulation of Ras protein signal transduction) and, (R-HSA-388396, GPCR downstream signaling) (Figure S3A). The *GPR18*, *GPR183*, *GPR20*, and *DENND3* candidate genes were selected (Table S9).

**Figure 5.**
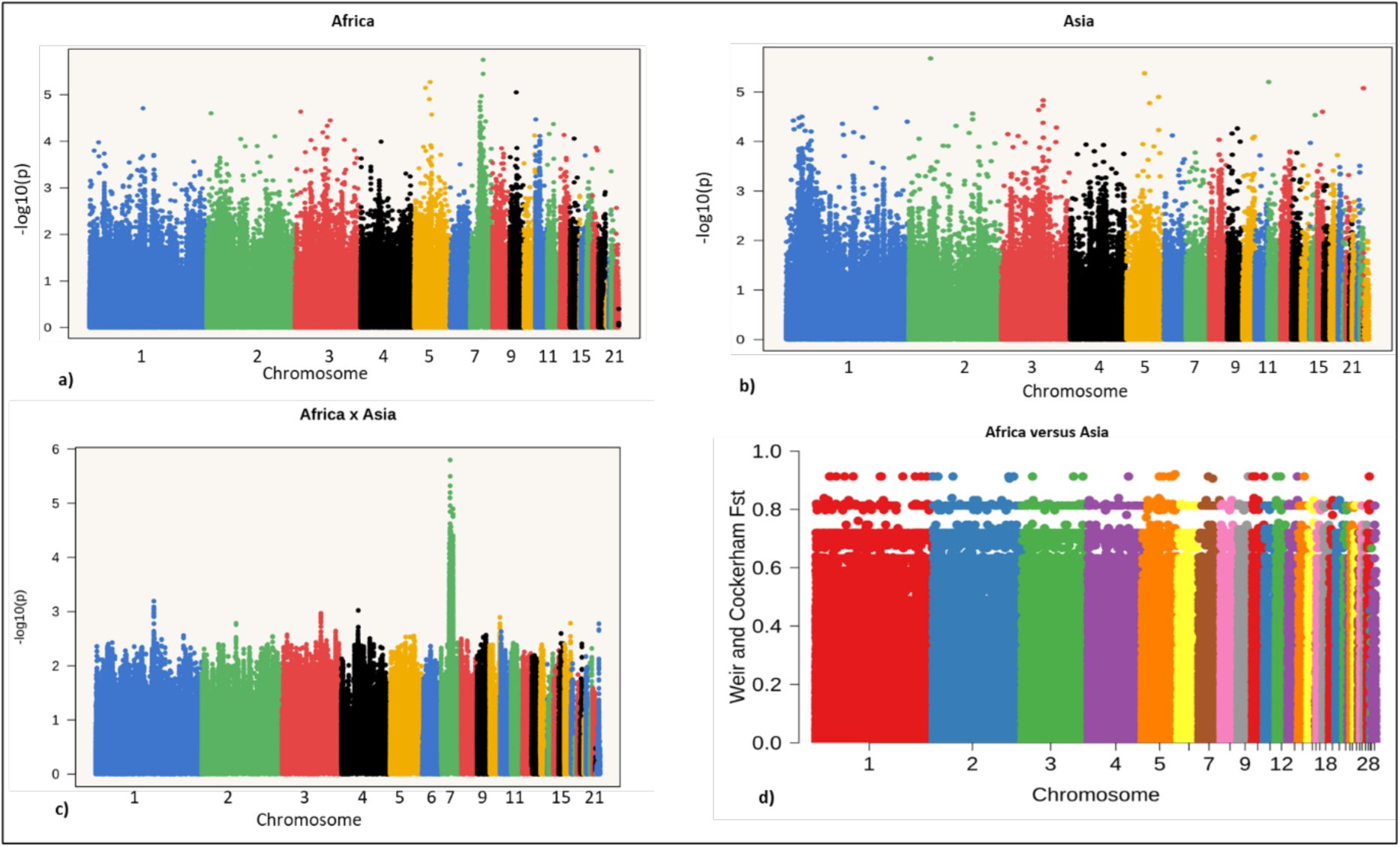
Genome-wide distribution of iHS, XP-EHH, and F*_ST_* scores for chicken populations from Africa and Asia. (a) Distribution of iHS values for chicken populations from Africa. (b) Distribution of iHS values for chicken populations from Asia. (c) Distribution of XP-EHH scores comparing amongst Africa vs. Asia chicken populations. (d) Distribution of F*_ST_* scores comparing amongst Africa vs. Asia chicken populations.

The Ras protein is responsible for activating a variety of signaling molecules by transporting them to the plasma membrane (Simanshu et al., 2017). This protein is vital for regulating cell growth, differentiation, apoptosis, cell migration, and neuronal activity (Olson & Marais, 2000; Simanshu et al., 2017). The GPCR (G-protein-coupled receptor) downstream signaling is important in mediating the cellular response to a wide variety of stimuli including hormones, neurotransmitters, and sensory signals (Guangxin et al., 2019). Given their crucial role in multiple signaling pathways involved in various cellular functions, the Ras protein, and the GPCR downstream signaling enriched terms may play a pivotal role in the adaptation of the chicken to their surroundings.

#### 3.4.2. Selection Signatures Common in Chicken Populations from Asia

The iHS analyses detected regions of putative selection signatures, notably with strong signals at chromosome 1 (GRCg6a position 34,551,571-34,761,125 bp), chromosome 2 (GRCg6a position 34,651,269-34,954,644 bp), chromosome 3 (GRCg6a position 34,777,526-34,790,199 bp), chromosome 4 (GRCg6a position 34,628,096-34,640,529 bp), and chromosome 7 (GRCg6a position 34,721,338-34,761,957 bp) (Figure 5(b)). One of the significantly enriched terms was the response to stimulus category (GO:0006979, response to oxidative stress) (Figure S3B). The response to oxidative stress has also been reported as one of the enriched terms in Chinese chickens inhabiting highland areas (Nan et al., 2023) The *HEATR5A*, *HNRNPU*, *KIF5C*, *PPP2CB*, *RBPMS*, *MGMT*, *EPC2*, *ADSS*, *WRN*, *TBC1D5*, and *GRIP1* were among the selected candidate genes (Table S10).

Oxidative stress occurs when there is an imbalance between the generation of reactive oxygen species (ROS) and the capacity of the cell to eliminate them (Ramiah et al., 2022; Rehman et al., 2018). ROS can harm DNA, proteins, and lipids, thereby leading to cellular impairment and possibly contributing to the onset of various illnesses. When chickens are exposed to both poor air quality and pathogens, it can lead to the generation of ROS (Nan et al., 2023). Hence, the response to the oxidative stress pathway could assist chickens in managing these stressors and protecting their cells from harm.

#### 3.4.3. Selection Signatures Common in Chicken Populations from Africa and Asia

Strong selection sweep regions were detected on chromosome 5 (GRCg6a position 5,444,151-5,461,448 bp), chromosome 14 (GRCg6a position 5,307,535-5,318,482 bp), chromosome 17 (GRCg6a position 5,345,787-5,353,651 bp), chromosome 20 (GRCg6a position 5,418,333-5,446,344 bp), chromosome 21 (GRCg6a position 5,586,241-5,605,153 bp), chromosome 26 (GRCg6a position 5,221,530-5,245,884 bp) and chromosome 27 (GRCg6a position 5,360,090-5,397,664 bp) by the XP-EHH method (Figure 5(c)) and on chromosome 1 (GRCg6a position 55,281,097-55,330,373 bp), and chromosome 2 (GRCg6a position 55,454,629-55,529,287 bp) by the F*_ST_* method (Figure 5(d)). One of the significant GO terms was the developmental process category (GO:0051960, regulation of nervous system development), (GO:0007420, brain development), and (GO:0043588, skin development) (Figure S3C). Some of the candidate genes that were identified included *ATP2B4*, *MAPT*, *CNBP*, *IGF-1*, *CLCN6*, *SLC12A7*, *SLC27A4*, *SREBF1*, *LRRC8A*, *HNF4A*, and *PAX6* (Table S11).

The enriched terms related to the regulation of nervous system development and brain development could enhance sensory perception, cognitive abilities, or behavioral traits that prove advantageous in the specific environment where the chickens inhabit. One of the genes that was identified in the metabolic process of transcription by RNA polymerase II is the cellular nucleic acid-binding protein (*CNBP*), also known as zinc-finger protein 9 (*ZNF9*).

*CNBP* is a highly conserved zinc-finger protein that has been associated with diverse cellular functions, including transcription and translation (Chen et al., 2018). Liang et al. (2008) highlighted the role of *CNBP* in the regulation of immunity and inflammatory responses. The gene may have an important function in supporting the immune system of chickens in heat-stressed environments.

Skin development was among the top enriched terms in the two chicken populations. The skin plays an important role in thermoregulation and acts as a protective barrier against diseases. The pigmentation can protect against harmful UV radiation (Henrique et al., 2023). The color of the skin can also impact the absorption of vitamin D, which is necessary for bone health (Akinyemi & Adewole, 2021). One of the annotated genes in the developmental process of the skin was the *COL1A1* gene, which codes for type I collagen, and is the main structural component of the extracellular matrix of the skin (You et al., 2023). This gene is essential for the development and maintenance of healthy skin since it provides tensile strength and elasticity to the tissue, attributes which are important for the protection of poultry from external harm.

## 4. Conclusions

The wild vulturine guineafowl species from Africa was shown to be of pure breed, as well as some indigenous chicken from Ethiopia-Jarso, and China-Yunnan. These two poultry species can be used for breeding practices.

Some of the selected candidate genes were specific to each of the analyzed guineafowl populations (Africa vs. Asia, Africa vs. Europe, Europe vs. Asia, Africa, Asia, Europe), and chicken populations (Africa vs. Asia, Africa, Asia), while some of the genes and pathways were shared across the analyzed populations. Among the shared genes between the guineafowls and chicken populations include the *ATP2B4* which was selected for the guineafowl populations from Africa vs. Europe and Chicken populations from Africa vs. Asia. The *CNBP*, *SREBF1*, *HNF4A*, *LRRC8A*, and *SLC27A4* genes were selected for the chicken populations from Africa vs. Asia, and guineafowl populations from Africa vs. Europe, and Asia. Most of the genes and pathways selected in this study have been reported to be involved in maintaining homeostasis and regulating stress, which are crucial factors for breeding poultry that can adapt to different climates.

## Supporting information

https://docs.google.com/spreadsheets/d/15rKN3UER_eKaURKjjPEHrIrdQJ8WvWjt/edit?gid=634045355#gid=634045355

## Data Availability

The supporting data are enclosed as additional files

## Supplementary Information

**Figure S1:** GO enrichment analysis for the candidate genes was revealed in pairwise comparison between guineafowls from Africa, Asia, and Europe

**Figure S2:** GO enrichment analysis for the candidate genes was revealed in a pairwise comparison between guineafowls from Africa, Asia, and Europe

**Figure S3:** GO enrichment analysis for the candidate genes was revealed in a pairwise comparison between chicken from Africa and Asia

**Table S1:** Summary of guineafowl samples from selected countries in Africa, Asia, and Europe

**Table S2:** Summary of chicken samples from selected countries in Africa and Asia

**Table S3:** Candidate genes in selected pathways and GO terms of relevance in guineafowl populations from Africa and Asia

**Table S4:** Candidate genes in selected pathways and GO terms of relevance in guineafowl populations from Africa and Europe

**Table S5:** Candidate genes in selected pathways and GO terms of relevance in guineafowl populations from Europe and Asia

**Table S6:** Candidate genes in selected pathways and GO terms of relevance in guineafowl populations from Africa

**Table S7:** Candidate genes in selected pathways and GO terms of relevance in guineafowl populations from Asia

**Table S8:** Candidate genes in selected pathways and GO terms of relevance in guineafowl populations from Europe

**Table S9:** Candidate genes in selected pathways and GO terms of relevance in chicken populations from Africa

**Table S10:** Candidate genes in selected pathways and GO terms of relevance in chicken populations from Asia

**Table S11:** Candidate genes in selected pathways and GO terms of relevance in chicken populations from Africa and Asia

